# Enhanced endogenous gene tagging in human induced pluripotent stem cells via AAV6-mediated donor delivery

**DOI:** 10.1101/2024.05.24.595765

**Authors:** Erik A. Ehlers, Kyle N. Klein, Margaret A. Fuqua, Julia R. Torvi, Javier Chávez, Lauren M. Kuo, Jacob McCarley, Jacqueline E. Smith, Gaea Turman, Danielle Yi, Ruwanthi N. Gunawardane, Brock Roberts

## Abstract

Systematically tagging endogenous proteins with fluorescent markers in human induced pluripotent stem cells (hiPSCs) allows observation of live cell dynamics in different cell states. However, the precise insertion of fluorescent proteins into live cells via CRISPR/Cas9-induced editing relies on homology-directed repair (HDR). The nonhomologous end-joining (NHEJ) DNA repair pathway often outcompetes HDR, resulting in irreversible insertions and deletions (INDELs) and low knock-in efficiency. Recognizing successful HDR-mediated tagging events is an additional challenge when the target gene is not expressed in stem cells and successful tagging cannot be immediately observed. To address these challenges, we used: 1) adeno-associated virus serotype 6 (AAV6) mediated DNA donors at optimized multiplicity of infection (MOI) to deliver tag payloads at maximal efficiency; 2) titrated, multiplexed Cas9:gRNA ribonucleoprotein (RNP) amounts to assure balanced HDR/INDEL frequency among conditions; 3) long-amplicon droplet digital PCR (ddPCR) to measure the frequency of HDR-generated alleles in edited pools; and 4) simultaneous Inference of CRISPR Edits (ICE) to detect and thereby avoid conditions significantly saturated (>50%) with INDELs. These approaches enabled us to identify efficient and accurate editing conditions and recover tagged cells, including cells tagged at loci not expressed in stem cells. Together these steps allowed us to develop an efficient methodology and workflow to clonally isolate directly from an ideal cell pool with optimal HDR and minimized INDEL frequencies. Using this approach, we achieved both monoallelic and biallelic insertion of fluorescent markers into four genes that are turned on during differentiation but not initially expressed in hiPSCs, where direct selection of tagged cells based on fluorescence was impossible: *TBR2, TBXT, CDH2* (pro-differentiation and pro-migratory genes), and *CDH5* (endothelial specific gene). Through a systematic evaluation of various gRNA sequences and RNP concentrations, we identified conditions for each gene that achieved high HDR frequencies, peaking at 38.6%, while also avoiding conditions saturated with INDELs, where isolation of clones with a tagged allele in trans with an unedited allele is difficult. Overall, this methodology enhances the efficiency of fluorescent tag knock-in at genes not expressed in hiPSCs, facilitating reliable image-based observation of cellular processes, and enables recovery of accurately edited mono- and biallelically tagged clones. We standardized these approaches to yield an efficient and general workflow for introducing large HDR mediated knock-ins into hiPSCs.

## Introduction

The development of CRISPR/Cas9 and related technologies have revolutionized cell biology because it enables site-specific integration of exogenous sequences at endogenous gene loci, a technology referred to as “endogenous tagging” (Bukhari and Müller, 2019). Endogenous tagging enables live-cell fluorescence imaging studies for understanding complex biological processes by tracking protein abundance, organization, and dynamics (Dambournet et al., 2018; Bukhari and Müller, 2019; Lau et al., 2020; Gerbin et al., 2021; Cho et al., 2022; Rafelski and Theriot, 2024; Viana et al., 2023). Several protocols have emerged to facilitate endogenous gene tagging at varying degrees of throughput and efficiency in mammalian cells. Those employing HDR exert the greatest control over the designed outcome but also the least efficiency, particularly in human pluripotent cells (Mikkelsen and Bak, 2023). Protocols to achieve efficient cell line tagging therefore remain an outstanding need.

Several challenges exist to producing precisely tagged hiPSC lines without prohibitive labor investment. First, despite improvements in the efficacy of HDR-based editing in hiPSCs, HDR remains a low-frequency outcome for both vector and single-stranded oligodeoxynucleotide (ssODN)-based donor strategies (Soldner et al., 2011; Zhang et al., 2017; Fehér et al., 2022). Consequently, producing an edited hiPSC line often requires a fluorescence or antibiotic selection scheme or labor-intensive clone screening to find rarities with the intended genetic alteration. We aimed to build upon recent successes using CRISPR/Cas9-mediated site-directed knock-ins using AAV6 DNA donors in human cells in hopes that selection of any kind would be ultimately unnecessary (Bak et al., 2017; Gao et al., 2019; Martin et al., 2019; Fu et al., 2021). These studies reported remarkable increases in HDR efficiency, with successful tagging at over 70% of total alleles, including in hiPSCs. MOI values used in these reports varied and we therefore sought to standardize them for use within the WTC-11 hiPSC system because this has been established as an important variable for HDR (Charlesworth et al., 2018).

Second, accurate tagging requires a strategy to moderate the deleterious contribution of INDELs introduced through NHEJ, particularly where monoallelic tagging is preferred to minimize biological consequences of the tag. To integrate fluorescent tags efficiently via HDR, RNP cleavage efficacy and targeting position relative to the intended site of tag insertion are crucial considerations (Boyle et al., 2021; Schubert et al., 2021). We reasoned that titrating the concentration of a series of different gRNA sequences targeting near the tag insertion site would reveal ideal experiment conditions for recovery of both monoallelic (“balanced” HDR and INDEL induction) and biallelic tagged (maximum HDR) clones. Accurate tagging also involves surveillance among tagged clones for inaccurate HDR, often responsible for complex sequence duplications and expansions, which has not been extensively reported on among AAV6-mediated tagging studies.

A third challenge involves tagging genes with undetectable expression in undifferentiated stem cells, which are often of interest because they influence differentiation-specific processes but are not expressed in the pluripotent state where editing takes place (Roberts et al., 2019). We reasoned that ddPCR detection of HDR alleles recombinant in the genome would function generally to identify ideal editing conditions after titrated RNP electroporation and AAV6 transduction. We report a generally applicable workflow using this method and employ INDEL surveillance as a simultaneous, complementary step for identifying conditions with optimized editing without saturating INDELs.

To address these challenges, we developed a workflow combining CRISPR/Cas9 with AAV6-mediated donor DNA delivery to enhance HDR efficiency in hiPSCs (Figure 1A). This methodology circumvents the limitations of traditional HDR techniques, which often result in low knock-in efficiencies. By integrating comprehensive surveillance tools such as ICE and custom ddPCR assays, we precisely quantified and optimized conditions for HDR, leading to improved genome editing outcomes. Our aim was to demonstrate this method’s feasibility by endogenously tagging the transcriptionally silent genes *CDH2, CDH5, TBR2*, and *TBXT* entirely via a PCR-driven strategy. Whenever feasible, the selection of protein terminus and linker for each tagging experiment was informed by insights from the literature and through direct consultations with researchers who had previous experience with tagging the target of interest (Kurokawa et al., 2017; Tosic et al., 2019; Sheppard et al., 2021). Here, we document the knock-in efficiencies, editing precision, and single-cell cloning outcomes, effectively creating isogenic cell lines at each targeted locus. This methodology offers a reliable and efficient strategy for endogenous protein tagging in hiPSCs, thereby expanding the potential for live-imaging studies of internal cellular organization and differentiation processes.

**Figure 1.**
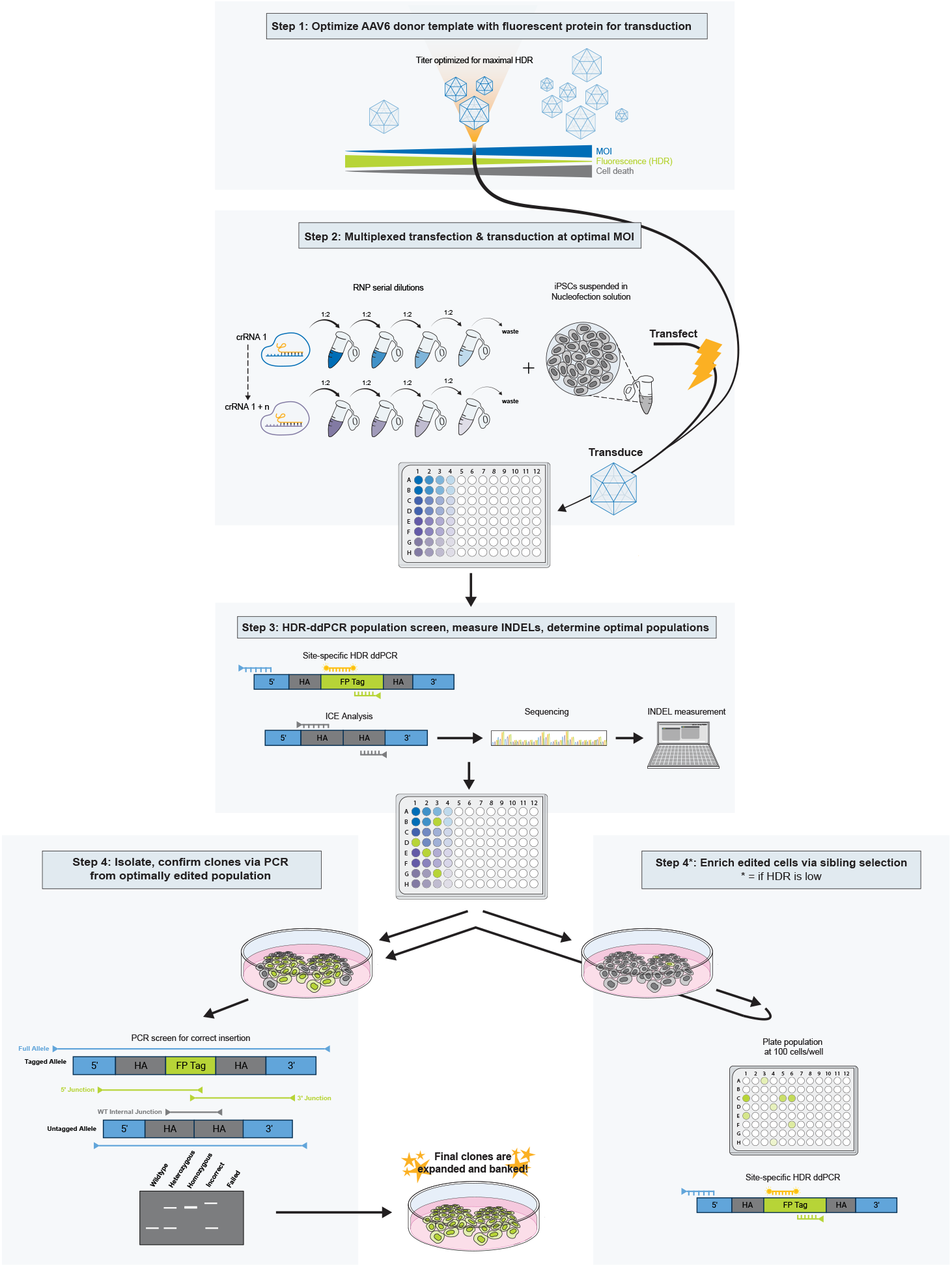
Graphical overview of enhanced endogenous gene tagging in human induced pluripotent stem cells. (**A**) Step 1: Optimize AAV6 MOI for transduction. Viral MOI is optimized to balance HDR and cell survival. A concentration of viral titer (blue wedge) is chosen based off the highest fluorescent protein expression (green wedge) for the lowest amount of cell death (gray wedge). Step 2: Multiplexed transfection and transduction at optimal MOI. The MOI determined in step 1 is used following RNP transfection to insert the donor template via HDR. Transfection conditions are multiplexed targeting multiple target sequences among multiple RNP dilutions (blue and purple tubes). Cells are then plated in a 96-well plate. Step 3: HDR-ddPCR population screen, measure INDELs, and determine optimal populations. Site-specific ddPCR is done to determine HDR-edited populations. Primers (blue and green) are used to amplify the edited gene of interest. ddPCR probes (orange) designed to the fluorescent protein tag give specific amplification of HDR-edited alleles. PCR (gray primers), sequencing, and ICE analysis is also done to determine INDEL rates. Based off the results of step 3, and the HDR-editing efficiency, two avenues can be pursued. If HDR is high, move on to step 4: isolate and confirm clones via PCR from the optimally edited population. A series of three different PCRs are run to assay for the full edited allele (blue arrow), the junction between the edit and the genome (green arrows), and within the RNP cut site (gray arrow). The resulting PCR can be visualized on a gel (bottom gray box) to screen for both insertions and copy number. This is followed up by Sanger sequencing to verify coding sequences are correct. Final clones can then be expanded and banked. If the result of step 3 was low levels of HDR, follow step 4*: enrich edited cells via sibling selection. Here cells are plated at 100 cells per well. This increases the frequency of finding a well with high HDR when assayed via ddPCR. Newly identified enriched populations can return to the workflow, following step 4, until final clones are expanded and banked.

## Results

### Optimizing MOI for AAV6-mediated tagging in WTC-11 hiPSCs

We employed AAV6 donor templates because of their demonstrated effective and specific DNA introduction through HDR in hiPSCs (Martin et al., 2019; Fu et al., 2021). We custom-ordered AAV6 donors with variable-length homology arms (HAs) corresponding to the centromeric protein *CENPA*, the mitochondrial transporter *TOMM20*, and the histone *HIST1H2BJ*. Donors encoded target-specific linkers, and either mEGFP or mTagRFP-T as tag sequences (Methods), features that were consistent with our previous tagging work using plasmid donors (Roberts et al., 2017, 2019). We chose initial gene targets (*CENPA, TOMM20*, and *HIST1H2BJ*) that are expressed in stem cells and that we had previously tagged using plasmid donors. Because all targets are expressed in stem cells, we could rapidly measure HDR using flow cytometry while optimizing the feasibility of AAV6 donor template deliveries. We performed all experiments with the WTC-11 hiPSC line with the eventual goal to expand the Allen Cell Collection (allencell.org) (Kreitzer et al., 2013).

We tested various MOIs to identify an optimum balance between cell viability and tagging efficiency and found >6% peak HDR among the three tagging experiments (Figure 2A-C). Furthermore, tagging efficiency >5% was observed at MOIs permitting >50% cell viability (Figure 2D-F). We used the percentage of fluorescent protein positive (FP+) cells to measure HDR efficiency four days after editing, while simultaneously using DAPI staining as a readout for cell viability. We observed that cell viability declined precipitously at increasing MOI, regardless of whether cells were electroporated. To quantify this, we used the percentage of live cells (DAPI-negative) four days after transfection/transduction normalized to live untreated control cells. MOI titration experiments on *CENPA* showed an optimal MOI range between 3,130-12,500 for transduction, with FP+ cell percentages ranging from 4.1-6.8% and cell survival rates of ∼25% (Figure 2A and D). For *TOMM20* MOI titration experiments showed an optimal range between 1,560-6,250 with FP+ cell percentages ranging from 4.1-7.8% at a cell survival rates between 25-50% (Figure 2B and E). For *HIST1H2BJ*, we observed no HDR efficiency peak. Instead, efficiency increased at higher MOI throughout the tested range, reaching the peak HDR of 10.2% at MOI 10,000 but with only 9.5% viability. Therefore, we chose the range of 3,130-12,500 MOI as the optimal range because it showed ∼50% cell survival along with 2.5-4.4% FP+ cells (Figure 2C and F), balancing efficient HDR and adequate cell survival to avoid extreme bottlenecking of cell population before downstream clonal isolation. From these results, we therefore performed subsequent transfections at an MOI of 4,500 because this value represented an approximate average inflection point in diminishing cell viability attributable to AAV6 transduction alone. This value was notably reduced compared to other reports, where 3x10^3^-3x10^5^ has been reported (Martin et al., 2019; Fu et al., 2021). We also noted that cell survival with AAV6 only declined at MOI values approximately ∼2-fold higher than ideal values for tagging. These data suggest that simple determination of the MOI responsible for ∼50% cell death after transduction likely suggests a near-optimum value for gene tagging in subsequent experiments.

**Figure 2.**
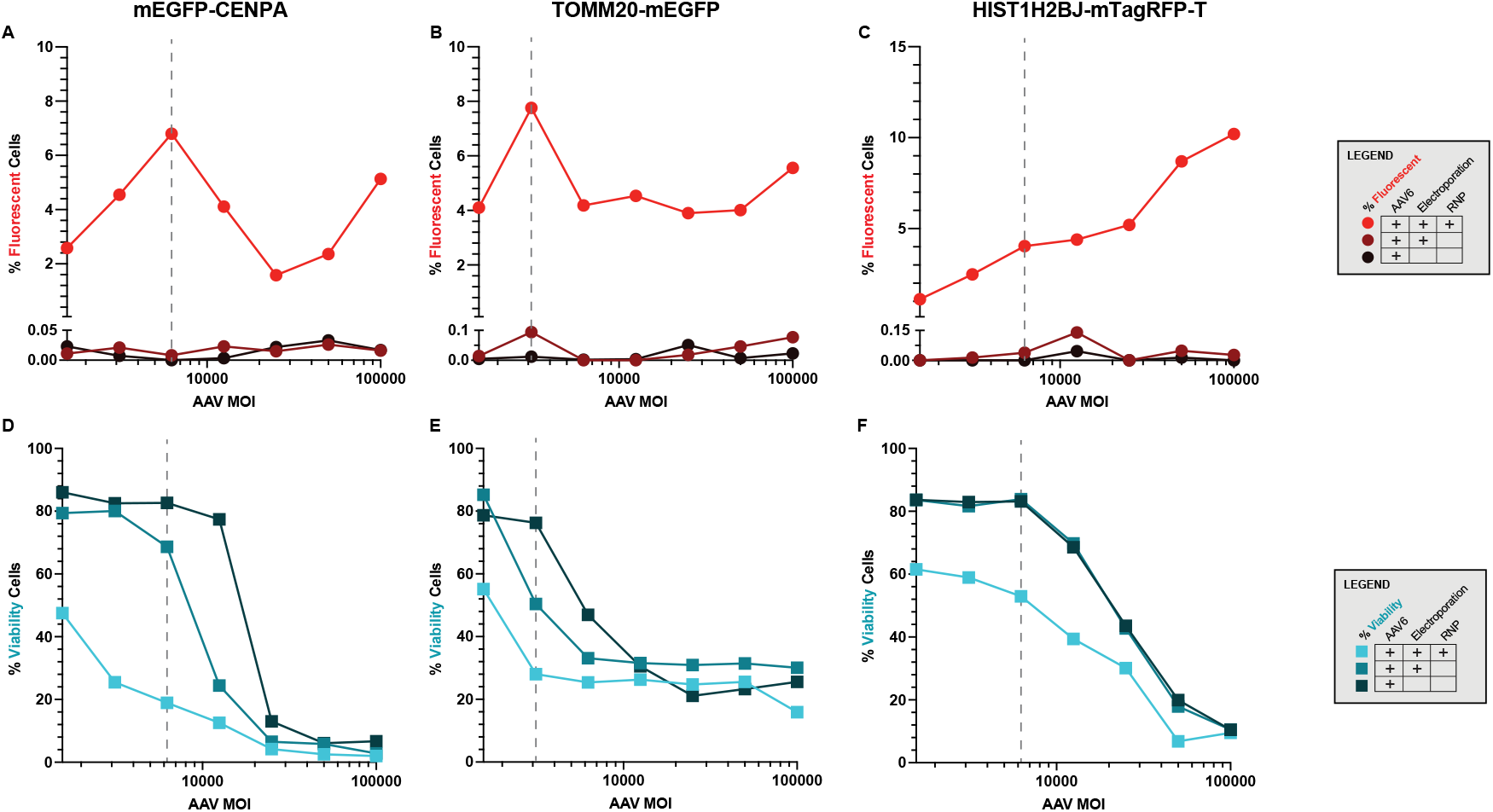
AAV6 MOI titration compared to fluorescent protein expression and cell survival. **(A-C)** Graphs show the percentage of cells fluorescent via FACS (red round data points) on the y-axis and AAV6 MOI on the x-axis, in logarithmic base 10 scale. **(D-F)** Graphs show percentage of viable cells using DAPI to discriminate live and dead cells via FACS (blue square data points) on the y-axis and AAV6 MOI on the x-axis, in logarithmic base 10 scale. **(A-F)** All graphs legends show the experimental condition, AAV+RNP, which includes both the AAV6 transduction and the RNP transfection (red or blue). Additionally, there are the controls, -RNP, which does not include the RNP in the transfection (dark red or blue), and -RNP -Electroporation, which is just the AAV6 transduction alone (black). Gray dashed line indicates the center point of the optimal MOI range. **(A and D)** When cells were edited for *mEGFP-CENPA*, the range of 3,130-12,500 MOI showed ∼20% cell survival along with the highest expression of fluorescent cells. **(B and E)** Cells edited with *TOMM20-mEGFP* showed the highest fluorescent protein expression and cell survival ∼25% in the range of 1,560-6,250 MOI. **(C and F)** *HIST1H2BJ* tagged with mTagRFP-T showed a different trend of fluorescent protein expression where the peak was not lower MOI ranges, but the expression increased with MOI. However, the range of 3,130-12,500 MOI, showed greater than 50% cell survival along with an acceptable percentage of cells fluorescing.

### Rapidly quantifying DNA repair outcomes among multi-plexed editing conditions

Relying on Fluorescence-activated Cell Sorting (FACS) to assess HDR is limited to expressed gene targets, prompting us to develop an alternative HDR surveillance assay for broader screening purposes. This assay overcomes FACS limitations by quantifying unique HDR alleles in populations within days of editing regardless of gene expression. To that end, we designed a ddPCR assay to quantify HDR allele frequency. For each edited locus, we positioned one primer distal to one homology arm and the other, along with the probe, within the tag sequence, ensuring only recombinant editing events are detected (Supplement Figure 1). Equally importantly, to minimize potential effects on endogenous processes due to editing, no change to the coding sequence of either allele is desired aside from installing the protein tag. An additional step to assess INDEL formation in addition to HDR is therefore crucial because INDELs may inactivate affected alleles (Miyaoka et al., 2016). To quantify INDELs in edited populations, we PCR-amplified the region encompassing the RNP target site and performed Sanger sequencing, followed by ICE analysis (Conant et al., 2022).

We envisioned that these assays would allow us to rapidly identify populations of edited hiPSCs with a favorable balance of HDR and INDELs. To assess this assay strategy, we investigated gene-tagging conditions among four genes silent in hiPSCs but activated after differentiation: *TBR2, TBXT, CDH2* and *CDH5*. To produce a dual-tagged line, we introduced the *CDH2-mTagRFP-T* donor into a previously generated CDH1-mEGFP line (allencell.org). We designed and individually tested 2-4 gRNA targeting sequences at each gene while titrating RNP concentrations from 5-40 μM. We were mindful that crRNA:tracrRNA duplexes (gRNAs) and synthetic modified single gRNAs (sgRNAs) with enhanced reported stability have differing reported nuclease activities. Evaluation on a case-by-case basis may indicate preferred chemistry for balancing the yield of HDR- and INDEL-modified alleles (Dang et al., 2015). Following nucleofection, cells were transduced with an AAV6 donor template containing homology arms specific to each gene, target-specific linkers, and mEGFP, mTagRFP-T or HaloTag as tags at an MOI of 4,500, chosen as a value representative of optimized conditions from initial trials (Figure 2). We then assayed RNP nuclease efficiency using ddPCR and used ICE to detect INDELs. For all experiments, we included a control gRNA targeting the safe harbor AAVS1 (adeno-associated virus site 1) as well as mock-electroporated and “untreated” negative controls. Synthetic DNA PCR template mixed with gDNA was included as a positive control (Methods).

Site-specific HDR predictably varied depending on the gRNA targeting sequence, RNP concentration, and the gene targeted. Higher RNP concentrations generally increased HDR (Figure 3A-D). Multiplexed transfection conditions and subsequent screening identified the targeting sequence and RNP concentration most efficient at site-specific HDR. HDR was maximized at 26.3%, 6.4% and 9.8% in initial tagging experiments targeting *CDH5, TBXT* and *CDH2*, respectively (Figure 3A-C). We re-tagged *TBXT* knowing that crRNA #5 most effectively targeted the locus for HDR delivery of mTagRFP-T (Figure 3B), substituted the crRNA:tracrRNA duplexed gRNA design used in initial targeting with a sgRNA, used an alternative HaloTag donor sequence and increased the RNP up to 140 μM. With these optimization strategies, HDR reached 38.7% (Figure 3D).

**Figure 3.**
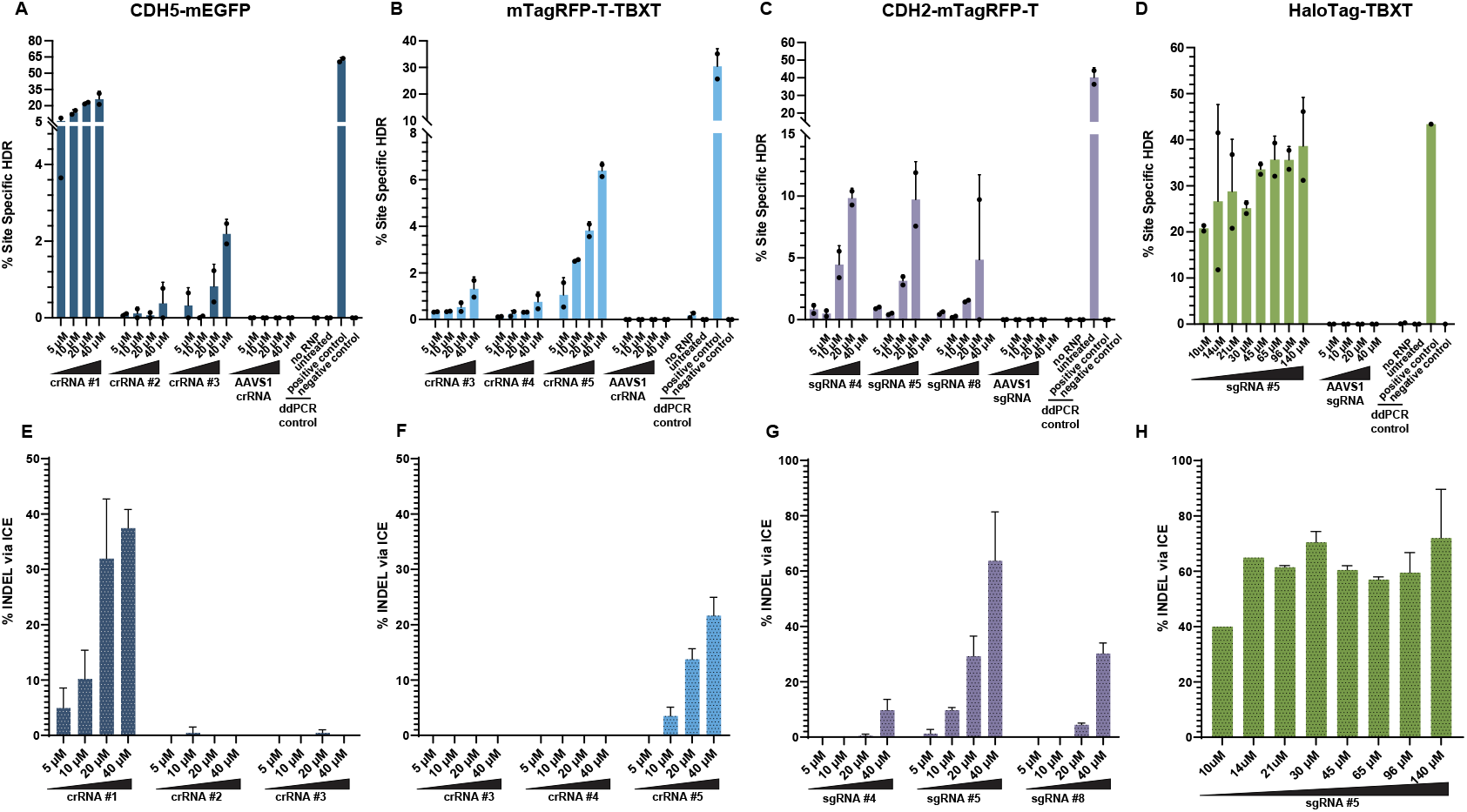
ddPCR detection of HDR and INDEL frequency via ICE among variable target sequences and RNP concentrations. **(A-D)** Different gRNAs tested across different RNP concentrations along with transfection controls (no RNP, untreated), and ddPCR controls are shown on the x-axis. AAVS1 gRNA was used as a negative control for HDR editing. Transfection control, no RNP, still electroporates the cells and contains all the reagents for transfection except the RNP complex. Control cells were cultured under the same conditions as the experimental samples until the stages of transfection and transduction. Subsequently, these controls were seeded into a 96-well plate for continued cultivation. Controls for ddPCR include the positive control (synthetic gBlock DNA template) and negative control (unedited AICS-0 cells). The y-axis shows the percent of site specific HDR insertion via ddPCR. **(E-H)** The percent of the population showing INDELs determined via ICE is shown on the y-axis. On the x-axis are the various gRNAs tested across a range of RNP concentrations. **(A and E)** *CDH5-mEGFP* cells showed the highest rate of HDR **(A)** and INDELs **(E)** with crRNA #1. These rates increased with increasing RNP concentration. **(B and F)** *mTagRFP-T-TBXT* cells showed the highest percentage of HDR **(B)** and INDELs **(F)** with crRNA #5 where rate also increased with RNP concentration. **(C and G)** In *CDH2-mTagRFP-T* cells both sgRNA #4 and #5 showed high rates of HDR **(C)** but sgRNA #4 showed the lowest rates of INDELs via ICE **(G)**. In all cases, rates increased with RNP concentration. **(D and H)** *HaloTag-TBXT* edited cells showed increasing rates of HDR with increasing RNP concentration **(D)** but INDEL rates via ICE did not strikingly change with concentration **(H)**.

ICE analysis of INDEL induction allowed us to monitor increased Cas9 activity at increasing RNP amounts. Across the initially tested range of RNP concentrations (5-40 μM), 5-37.5%, 3.5-21.6%, and 1.3-63.7% INDELs were observed when targeting *CDH5, TBXT* and *CDH2*, respectively (Figure 3E-H). Re-editing *TBXT* with HaloTag using 10-140 μM RNP achieved 40-72% INDELs (Figure 3H).

We also employed sgRNAs for targeting *CDH2* with mTagRFP-T, encouraged by the elevated editing observed with *HaloTag-TBXT*. Notably, higher RNP concentrations typically led to an enhanced HDR percentage. At 40 μM RNP, we identified targeting sequences that yielded HDR efficiencies of 9.8%, 9.7%, and 4.9% (Figure 3C). Although these populations displayed similar HDR efficiencies, ICE analysis of *CDH2-mTagRFP-T* showed varied INDEL formation among the targeting sequences (Figure 3G). Specifically, sgRNA #4 at 40 μM resulted in populations with 9.8% INDELs, sgRNA #5 at 40 μM led to 63.8% INDELs, and sgRNA #8 at 40 μM induced 30.3% INDELs. These data underscore the advantage of screening multiple gRNAs at different RNP concentrations for HDR and INDELs because conditions with optimal HDR and relatively few INDELs can be selected for further monoclonal isolation.

### Genetic analysis of editing outcome precision in clones

To evaluate HDR and NHEJ precision, we generated clones through low-density plating and manual isolation from populations with optimally balanced HDR and INDEL induction, followed by PCR screening (Figure 4A-E). We applied PCR-based screening to distinguish between precise and imprecise HDR edits at the targeted locus (Roberts et al., 2019; Suchy et al., 2024). First, we performed PCR screening with primers distal to both donor homology arms (“full allele” PCR) (Figure 4A, blue arrows). We thus quickly determined individual clones’ allelic composition because mono- and biallelic edits show unique PCR patterns (Figure 4B). We then performed junctional PCR on clones confirmed by full allele PCR to ensure precise template incorporation at 5’ and 3’ junctions (Figure 4A, orange arrows). Finally, we performed PCR and Sanger sequencing to identify INDELs in the tagged and/or untagged allele (Figure 4A, gray arrows).

**Figure 4.**
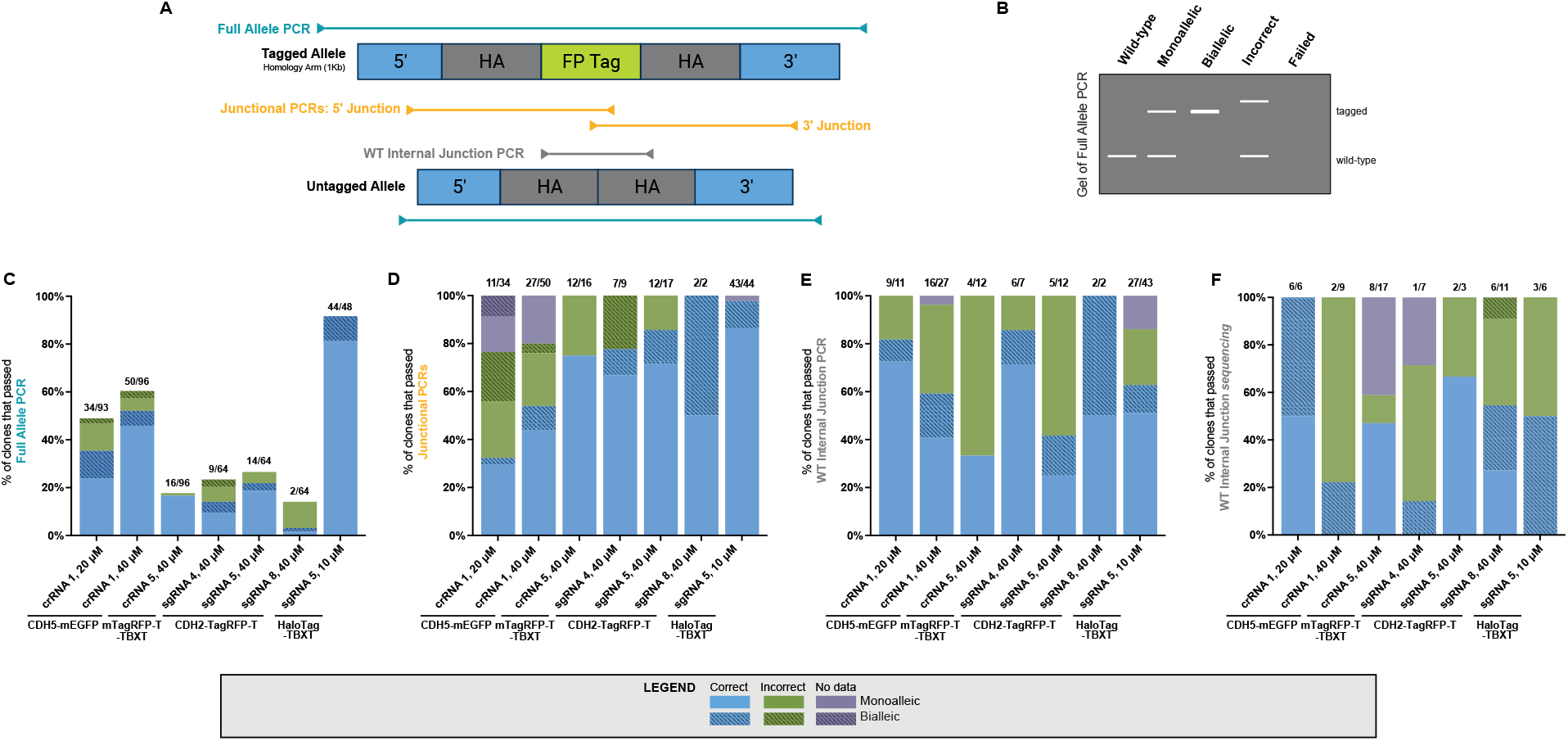
Surveillance of HDR-edited clones via PCR. **(A)** Schematic of the three steps of PCR surveillance. Full allele PCR (blue arrows) is where the region outside the homology arms are amplified. Junctional PCRs (orange arrows) are where the region outside the homology arms to the fluorescent protein tag is amplified. Wild-type (WT) internal junction PCR (gray arrows) is where the region inside the homology arm spanning the CRISPR cut site is amplified. **(B)** Schematic of the possible outcomes of PCR when visualizing via agarose gel. Unedited and tagged alleles are labeled on the right. **(C-E)** When we surveilled the four edited genes using full allele PCR **(C)**, junctional PCRs **(D)**, WT internal junction PCR **(E)**, and WT internal junction Sanger sequencing **(F)** clones are classified as either correct (blue), incorrect (green), or no data (purple). Edits that are “correct” indicate they had the correct allele size and sequence. Edits that are “incorrect” indicate that editing did occur, but the allele size and/or sequence is wrong. Lastly, edits that “failed” indicate that nothing was amplified. Clones are also grouped by monoallelic (solid color) or biallelic (striped color) edits. Sample numbers for each gene, gRNA, and RNP concentration tested are written above the bar, showing how many clones passed over the total number of clones surveilled at that step.

As an example, with *CDH5-mEGFP* (crRNA #1, 40 μM RNP) transfected clones, full allele PCR showed 52.1% (50 of 96) of clones had mEGFP insertions of the correct size (Figure 4C). Of 50 tagged clones, 45.8% (44/96) had monoallelic insertions and 6.3% (6/96) had biallelic insertions. Of the clones confirmed by full allele PCR, 54% (27 of 50) of *CDH5-mEGFP* (crRNA #1, 40 μM RNP) clones were also confirmed by junctional PCR screening (Figure 4D). Specifically, 50% (22/44) of putatively monoallelic clones had correct junctional sequences and 83.3% (5/6) of putatively biallelic clones had correct junctional sequences (Figure 4D). Of the clones confirmed by junctional PCR, 59.0% (16/27) of *CDH5-mEGFP* (crRNA #1, 40 μM RNP) clones were confirmed by wild-type (WT) internal junction PCR (Figure 4E). These data include 50% (11/22) of putatively monoallelic clones and all (5/5) putatively biallelic clones (Figure 4E). Specifically, 2 out of 9 were sequenced and determined to be INDEL-free around the targeted cut site (Figure 4F). Clones interpreted as biallelic were further confirmed via ddPCR to have two tag copies to rule out hemizygosity (Supplement Figure 2), which has been observed as a possible editing outcome and exists as a plausible alternative explanation when the untagged allele is not observed after PCR amplification (Simkin et al., 2022). The genomic copy number of the tag sequence was consistent with each clone’s interpretation as a mono- or biallelically edited tagging outcome (Supplement Figure 2). These data suggested absence of the reportedly common concatemerized donor delivery outcome and that clones were accurately tagged (Suchy et al., 2024).

### Strategies to identify rare alleles

We also desired a fluorescent fusion tagged cell line labeling the pro-differentiation transcription factor *TBR2* (EOMES) (Tosic et al., 2019). However, initial efforts with several duplexed crRNAs produced HDR below 0.1% at all RNP concentrations (Supplement Figure 3) suggesting that these RNP complexes introduced targeted double strand breaks inefficiently. Repeating with sgRNAs, we identified sgRNA #3 as the most efficient targeting sequence in our panel, but with HDR frequencies ranging only from 0.6% to 3.1% (Figure 5A), and INDELs ranging from 0.2% to 0.7% among tested populations (Figure 5B).

**Figure 5.**
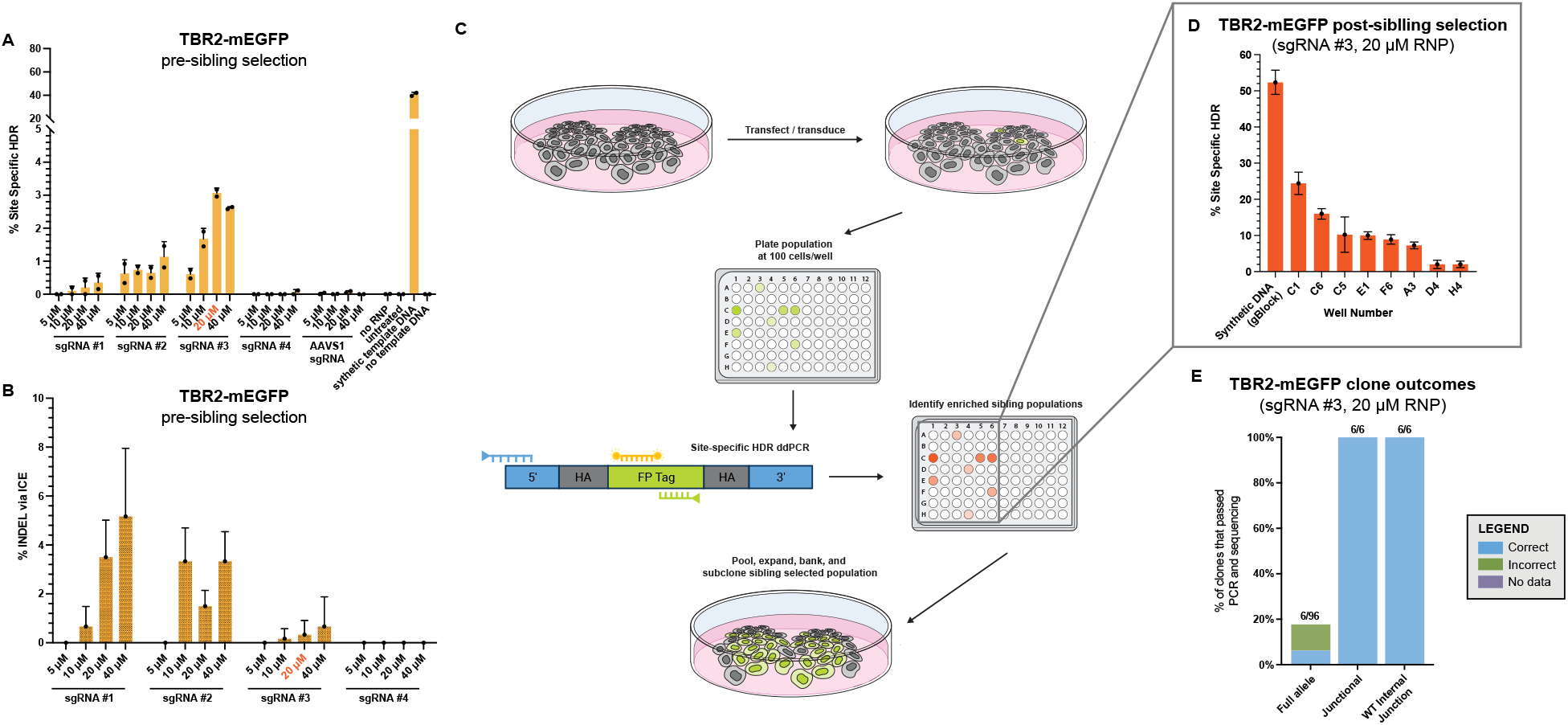
Sibling selection when HDR rates are low to assist in isolating correctly edited clonal populations. **(A)** *TBR2-mEGFP* pre-sibling selection showed low rates of HDR. In general, HDR rates increased with increasing RNP concentration with sgRNA #3 at 20 μM RNP (red text) showing the highest HDR percentages. **(B)** Similarly, INDEL rates via ICE showed the lowest rates with sgRNA #3 at 20 μM RNP (red text). **(C)** Overview of the sibling selection methodology. Cells are first transfected and transduced. Cells are then plated at low density, 100 cells/well, in a 96-well plate. Site specific ddPCR is then done on each population to identify wells with high rates of HDR. These wells are then expanded, subcloned, validated via PCR, expanded, and banked. **(D)** Post-sibling selection, the percentage of HDR increased. Well numbers of colonies tested are on the x-axis along with the positive control, synthetic gBlock DNA. **(E)** Post-sibling selection clone outcomes are graphed through the steps of PCR validation. On the x-axis is the step of PCR validation done, and on the y-axis is the percentage of clones passing that PCR and sequencing step. Atop the bars are the number of clones that passed that step over the total amount assayed.

To mitigate the low frequency of presumptively tagged cells in this cell pool, we attempted to enrich the edited allele frequency in this population before monoclonal isolation via sibling selection, a technique frequently utilized in yeast genetics for isolating a specific rare cell type by repeatedly subdividing a population of cells to enrich and isolate cells containing a rare allele, and its usage in hiPSCs for enriching single-base changes has been documented (Miyaoka et al., 2014). We adapted this process using HDR-ddPCR to identify edited subpools with the highest HDR frequency, after low-density plating.

To perform sibling selection, we plated the 20 μM sgRNA #3 *TBR2* population in a 96-well plate format with 100 cells per well (yielding on average an estimated 10-20 founding colonies) to enrich individual wells with an overall higher HDR percentage than the parental population. We then screened the 96 individual populations with the same long amplicon HDR-ddPCR as in the initial screen (Methods). We identified eight sibling-selected populations with HDR percentages over 5% from one 96-well plate, with three populations having 10-20% site-specific HDR (Figure 5D). To maintain outcome diversity within the enriched pool, we combined several selected sub-populations and proceeded with clonal isolation.

Full allele PCR of clones from the sibling-selected *TBR2-mEGFP* population revealed 6.3% (6/96) of clones with correctly sized mEGFP insertions (Figure 5E). Of the six clones observed with a correctly tagged edit, all were biallelic insertions. Clones validated with full allele PCR were re-analyzed with junctional PCR to confirm correct HDR. We then performed Sanger sequencing with primers positioned near the RNP cleavage site to confirm absence of wild-type bands and to identify potential INDELs, with all clones again confirmed. This comprehensive approach, culminating in the successful identification and validation of precisely edited biallelic clones without INDELs at the targeted locus, underscores the efficacy of sibling selection and PCR screening strategies for recovery of cells with rare alleles.

In summary, we 1) validated findings of increased HDR efficiency using AAV6-mediated donor sequences within hiPSCs (Martin et al., 2019; Fu et al., 2021); 2) varied RNP conditions in multiplex to identify optimal editing parameters; 3) screened editing conditions with efficient and high-throughput PCR-based methods to detect optimized HDR and INDEL allele frequencies and; 4) used these methods to successfully tag the differentiation-specific genes *CDH5, CDH2, TBR2*, and *TBXT*.

## Discussion

AAV6 vectors have emerged as a method to trigger efficient HDR for site-specific genome engineering in several systems, including hiPSCs. This report examines HDR delivery efficiency of fluorescent protein tag-sized payloads to seven loci, suggesting that efficient tagging within hiPSCs, including biallelic edits (provided cells are viable) may be generally feasible if appropriate targeted RNP activity can be achieved. We find that with this method >5% tagging allows us to identify >5 tagged candidate clones by screening only ∼10^2^ clones, and we achieved that level of efficiency in this report. Tagging efficiency may generally exceed the logistical minimum to avoid laborious, expensive, low-throughput enrichment techniques including FACS and antibiotic selection. Beyond making tagging experiments fundamentally feasible, avoiding antibiotic selection and FACS may also reduce stress experienced by cells as they undergo editing. Also, AAV6 donors trigger significant cell death after transduction, but only ∼2-fold greater in transduced electroporated cells as compared to transduced only. These further reduce barriers to tagging because unelectroporated cells can be subjected to a simple MOI titration after AAV6 acquisition, with the MOI permitting ∼50% normal growth likely optimal for tagging and related large-payload HDR-mediated genomic installations.

We developed a general PCR-based method to identify and measure HDR-edited allele frequency, providing added rationale to avoid alternative selection methods beyond elevated AAV6-mediated HDR efficiency. Long-amplicon ddPCR underpinned this approach, which has also recently been employed to detect rare events with clinical significance (Lasham et al., 2020) and to detect large payloads delivered by HDR (Martin et al., 2019). This strategy negated the need for a selection scheme due to the gene being expressed only after hiPSC differentiation and suggests a more streamlined workflow than our previous report (Roberts et al., 2019) for tagging genes specifically expressed during differentiation. We successfully applied this method for tagging four differentiation-specific genes, lending confidence that it may be a generally feasible approach to decoupling the tagging strategy from the gene’s expression state. We additionally demonstrated that this method could be used to enrich the frequency of tagged alleles in cell pools to make *TBR2-mEGFP* clonal isolation more tractable using a sibling selection strategy.

We also describe multiplexed titration in parallel of RNP complexes among dozens of potentially successful editing conditions. This approach is impactful because RNP activity specific to a given gRNA and its consequences on DNA repair is difficult to predict a priori. Algorithms predicting gRNA-specific RNP activity in terms of DNA double strand break induction (Doench et al., 2016) and likelihood of resulting in a deleterious INDEL (Bae et al., 2014) have been developed but may not apply to specific cellular contexts such as hiPSCs. The propensity of a gRNA to trigger HDR with a full-length fluorescent tag template remains unexplored by predictive algorithm development. It is therefore a sound strategy to evaluate many conditions in parallel and invest further effort in the optimally edited population, as described in this report. The ability to recover both monoallelic and biallelic clones is pivotal for live-cell 3D imaging studies, where the distinction between biallelic and monoallelic edits can have significant biological implications and where the two outcomes may be differently interpreted. This report suggests that biallelic edits may be found among editing conditions with maximal HDR where INDELs in trans among untagged alleles are irrelevant. By contrast clones tagged at one allele may be most readily derived from conditions with maximal HDR without exceeding our suggested threshold of ∼50% INDELs in the population. Differing INDEL/HDR induction with sgRNAs and crRNAs suggests gRNA chemistry as another variable to be exploited for identifying the optimal editing condition and a multiplexed experiment.

This gene tagging strategy yielded fluorescent endogenously tagged reporter lines for the epithelial-to-mesenchymal transition (EMT) (*CDH2, TBXT*, and *TBR2*) and endothelial cell differentiation (*CDH5*). Both differentiation processes represent dynamic changes in the transcriptome, cellular organization, and cellular behavior, making these processes attractive for investigation via 3D live cell microscopy. These EMT lines allow for real-time tracking of the EMT process, providing insight into the dynamics of this transition. These reporter lines can also enable lineage tracing and fate mapping of cells during early development, which is important for identifying regulatory pathways that control cell differentiation and state transitions (Tosic et al., 2019; Sheppard et al., 2021; Rafelski and Theriot, 2024). The *CDH5-mEGFP* reporter line enables direct observation of endothelial cells as they differentiate and is critical for studying the role of VE-cadherin in cell-cell adhesion, vascular structure formation, and endothelial layer integrity. Insights gained from this system may clarify mechanisms of vascular disorders characterized by disrupted cell adhesion (Kurokawa et al., 2017).

When creating fluorescently labeled hiPSC lines, we advise using both HDR and NHEJ surveillance assays to refine genome-editing efforts, particularly when NHEJ is an unwanted outcome. For genome editing with high precision, it is crucial to maximize HDR while minimizing NHEJ. Our technique evaluates both HDR and NHEJ prior to single-cell cloning, offering sensitivity and consistency. The preference for HDR over NHEJ varies across different genomic locations, indicating a complex genomic landscape that may involve epigenetic factors and DNA repair pathways. This method is poised to increase the ability to generate fluorescently tagged cell lines via DNA repair processes triggered by Cas9 nucleases, potentially leading to more efficient and precise genome-editing protocol. The approach we outline permits the multiplexed testing of genome-editing conditions, enabling the quantification of HDR and NHEJ while selecting against clones that have undergone NHEJ. If, however, NHEJ is the goal ICE can be employed to identify cell lines with NHEJ-induced gene disruptions (Conant et al., 2022). This assay allows for the monitoring of NHEJ frequency, and one can enrich for these events using sibling selection, as demonstrated by (Miyaoka et al., 2016). Therefore, this combined HDR and NHEJ protocol is an effective method for generating cell lines. Other available methods take advantage of the NHEJ repair pathway to introduce fluorescent tags (Artegiani et al., 2020) and these methods would also benefit from implementation of the strategy presented here in much the same way as it was used for HDR.

Despite the efficacy of this assay system, continual enhancements are necessary to keep up with the fast-evolving field. Additional methods, such as applying HDR-promoting drug combinations (Riesenberg et al., 2023) or inducing cold shock (Skarnes et al., 2019), might be necessary when sgRNAs and sibling selection alone are insufficient to achieve populations with a site-specific HDR frequency above 5%. With the ongoing advancements in DNA sequencing, ddPCR technology, and genome editing, this field is expected to stay at the forefront of scientific innovation. Presently, the combination of long-amplicon ddPCR and NHEJ monitoring stands as a reliable, swift, and cost-effective technique for evaluating genome-editing outcomes.

## Methods

### hiPSC tissue culture

All work with WTC-11 hiPSC cell line was approved by internal oversight committees and performed in accordance with applicable National Institutes of Health, National Academy of Sciences, and International Stem Society for Stem Cell Research Guidelines. WTC-11 was generated by the Bruce R. Conklin Laboratory at the Gladstone Institute and University of California-San Francisco (UCSF) and maintained using described methods (Kreitzer et al., 2013). The original cells received were at passage 33, and all passage numbers indicated in this study reflect additional subsequent passages.

The culture conditions used in this study have been described elsewhere (Roberts et al., 2017). Briefly, WTC-11 hiPSCs were cultured in a feeder-free system on tissue culture dishes or plates coated with Matrigel (Corning Inc., Corning, NY, USA) diluted 1:30 in cold DMEM/F12 (Thermo Fisher Scientific, Waltham, MA, USA). Undifferentiated cells were maintained in mTeSR1 medium (STEMCELL Technologies, Vancouver, BC, Canada) supplemented with 1% (vol/vol) penicillin-streptomycin (P/S) (catalog: 15140-122, Thermo Fisher Scientific, Waltham, MA, USA). Cells were not allowed to reach confluency greater than 85% and were passaged every 3-4 days by dissociation into single-cell suspension using StemPro Accutase (Thermo Fisher Scientific, Waltham, MA, USA). When in single-cell suspension, cells were counted using a Vi-CELL-XR Series Cell Viability Analyzer (Beckman Coulter, San Jose, CA, USA). After passaging, cells were replated in mTeSR1 supplemented with 1% P/S and 10 μM ROCK inhibitor (Y-27632, STEMCELL Technologies, Vancouver, BC, Canada) for 24 h. Medium was replenished with fresh mTeSR1 medium supplemented with 1% P/S daily. Cells were maintained at 37 °C and 5% CO_2_. A detailed protocol can be found at allen.org/written-protocols.html (Allen Institute for Cell Science, 2017).

### Clonal cell line generation

WTC-11 clonal cell line generation via limited dilution and manual isolation were updated from previously described methods (Roberts et al., 2017). Briefly, recently edited populations were seeded at a density of 1 x 10^4^ cells in a 10-cm Matrigel-coated tissue culture plate. After 5-7 days, clones were manually picked with a pipette and transferred into individual wells of 96-well Matrigel-coated tissue culture plates with mTeSR1 supplemented with 1% P/S and 10 μM ROCK inhibitor for 24 h. After 3-4 days of normal maintenance in mTeSR1 supplemented with 1% P/S, colonies were dispersed with Accutase and transferred into a fresh Matrigel-coated 96-well plate. After colony dispersion, cells were passaged 1:10 to a new 96-well Matrigel-coated plate for gDNA extraction with 20 μL of Quick Extract gDNA lysis solution (Catalog: QE09050, Bio-Search Technologies, VWR, Radnor, PA, USA). The remaining cells were overlaid with 50 μL of 20% mTeSR1, 60% KSR (Thermo Fisher Scientific, Waltham, MA, USA), 20% DMSO (SigmaAldrich, Inc., St. Louis, MO, USA) and transferred to 96 well-arrayed 300 μL capacity Cryo.S vials (Catalog: 976561, Greiner Bio-One, Monroe, NC, USA) with aid of a vial de-capping tool (Catalog: 852070, Greiner Bio-One, Monroe, NC, USA). Vials were transferred to −80 °C within 10 min for 1-3 days, then transferred to a vapor phase dewar for long-term storage. Cells generated in this study are listed in Figure 6.

**Figure 6.**
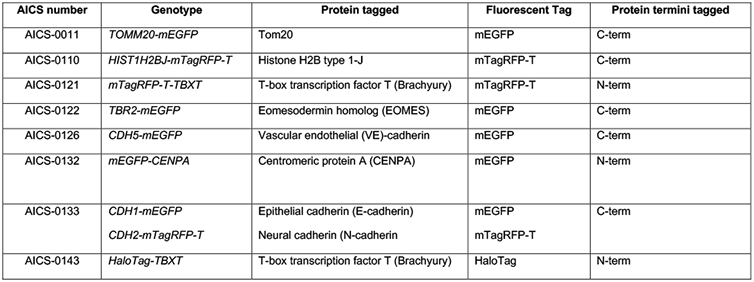
hiPSC lines generated in the study.

### AAV6 donor construction

AAV6 donor templates were designed uniquely for each target locus, with each following a similar design strategy. Homology arms 5’ and 3’ of the desired insertion site were each 1 kb in length and designed using the GRCh38 reference genome. WTC-11 specific variants identified from publicly available exome data (UCSC Genome Browser, n.d.). In cases in which the WTC-specific variant was monoallelic, the reference genome variant was used in the AAV6 donor template; when the WTC-specific variant was biallelic, the WTC-specific variant was used in the AAV6 donor template. When applicable, linkers for each protein were used to join the terminus of the protein with the fluorescent protein-encoding sequence (inserted 5’ of mEGFP for C-terminal tags and 3’ of mEGFP for N-terminal tags). AAV6 in-silico templates were provided to VectorBuilder (Chicago, IL, USA) where they were constructed and packaged. To prevent gRNAs from targeting the AAV6 donor template, we introduced mutations to disrupt RNP binding; when possible, these changes did not affect the amino acid sequence. Vector IDs can be used to retrieve detailed information about the vector on vectorbuilder.com *(CDH5-mEGFP: VB220217-1045xkx, CDH2-mTagRFP-T: VB210205-1204eyw, TBXT-HaloTag: VB220804-1195zvb, TBXT-mTagRFP-T: VB220214-1381hsh, TBR2-mEGFP: VB220329-1440cbm, CDH1-mEGFP: VB210205-1203yps, TOMM20-mEGFP: VB210921-1124vtp, H2B-mTagRFP-T: VB201020-1510wec)*.

### sgRNAs, crRNAs and Cas9 protein

To develop targeting sequences that facilitate double strand breaks near the desired genomic insertion site for the tag, we selected gRNA sequences based on their predicted effectiveness and specificity as described previously (Roberts et al., 2017). Briefly, we designed 3-5 targeting sequence per target locus, ensuring that the distance between the cut site and the desired fluorescent protein insertion site was less than 50 nucleotides, preferably under 10 nucleotides. Custom synthetic crRNAs and their complementary tracrRNAs were procured from IDT (Coralville, IA, USA) or Dharmacon (Lafayette, CO, USA). In cases where crRNA and tracrRNA combinations did not yield sufficient HDR for single-cell cloning, we utilized sgRNAs from Synthego (Redwood City, CA). These sgRNAs include 2’-O-methyl (OMe) analogs on the first and last three bases and 3’ phosphorothioate (PS) internucleotide linkages between the first three and last two bases. Both crRNA:tracrRNA duplexes and sgRNAs were reconstituted to 100 μM, and complexed with Cas9 protein in a 1:1 molar ratio. The recombinant wild-type *Streptococcus pyogenes* Cas9 protein was sourced from the University of California, Berkeley QB3 Macrolab (Berkeley, CA, USA). The targeted sequences with the PAM sequences highlighted are listed below:

~~~
*CENPA* (crRNA):
 5’- GGGCCCATGACACGCCGCAG**AGG**-3’
*CDH2* (sgRNA):
 sgRNA #4: 5’- GGAGGTGATGACTGAACTTC**AGG**-3’
 sgRNA #5: 5’- CTTGCTGACATGTATGGTGG**AGG**-3’
 sgRNA #8: 5’- TCAGCAAGTTTCTTGAACCG**TGG**-3’
*CDH5* (crRNA):
 crRNA #1: 5’- GGAGGAGCTGCTGTATTAGG**CGG**-3’
 crRNA #2: 5’- GCTGCTGTATTAGGCGGCCG**AGG**-3’
 crRNA #3: 5’- TAGGCGGCCGAGGTCACTCT**GGG**-3’
*HIST1H2BJ* (crRNA):
 5’ - ACTCACTGTTTACTTAGCGC**TGG** -3’
*TOMM20* (crRNA):
 5’ - AATTGTAAGTGCTCAGAGC**TGG** -3’
*TBR2* (sgRNA and/or crRNA):
 gRNA #1: 5’- GACACCTCAAAAGGCATGGG**AGG** - 3’
 gRNA #2: 5’ - GAGGTTAAAATAACTCTTTA**GGG** -3’
 gRNA #3: 5’ - TGAGGTTAAAATAACTCTTT**AGG** -3’
 gRNA #4: 5’ - ATTAGCTAACTTTTTGCAGA**TGG** -3’
*TBXT* (sgRNA and/or crRNA):
 gRNA #3: 5’- GGGGAGCTCATCCTCCCGTC**CGG** -3’
 gRNA #4: 5’- TTTCCCGCGCTCTCGGTGCC**AGG** -3’
 gRNA #5: 5’- TCCCGCGCTCTCGGTGCCAG**GGG** -3’
~~~

### Lonza nucleofection, AAV6 transduction, and gDNA extraction

To test multiple transfection conditions, we used the multi-well Lonza nucleofection system (4D Nucleofector, Lonza, Walkersville, MD, USA). hiPSCs were treated with 10 μM ROCK inhibitor 12-24 h in advance. When cells reached 70%-80% confluence, they were collected using Accutase (Thermo Fisher Scientific, Waltham, MA, USA). A nucleofection mixture was prepared by combining varying concentrations of Cas9 (40 μM, 20 μM, 10 μM, 5 μM) from the University of California, Berkeley QB3 Macrolab (Berkeley, CA, USA) with either crRNA:tracrRNA or sgRNA, each also at concentrations of 40 μM, 20 μM, 10 μM, 5 μM, maintaining a 1:1 molar ratio. This mixture was incubated for 10 min-at room temperature and further diluted with 20 μL of P3 Primary Cell solution (Lonza, Walkersville, MD, USA). For the nucleofection process, 3 x 10^5^ cells were combined with this nucleofection solution. The nucleofection was conducted using a 16-well Nucleocuvette Strip and a 4D Nucleofector system (Lonza, Walkersville, MD, USA), employing the CA137 electroporation code. Post-electroporation, the cells were immediately placed in a Matrigel-coated 96-well plate containing 200 μL of mTeSR media and 10 μM ROCK. The AAV6 donor vector was then introduced to the cells at a MOI of 3x10^3^-4.5x10^3^ and left to incubate at 37 °C for 24 h-. The media was replaced after 24 h-of editing, and the ROCK inhibitor was removed 48 h-later. Cells were cultured until reaching a confluency of 70-80% and then passaged 1:10 into Matrigel-treated 96-well plates. The remainder of the cells were spun and resuspended in 20 μL of Quick Extract gDNA lysis solution (Catalog: QE09050, Bio-Search Technologies, VWR, Radnor, PA, USA) and prepared for analysis. Passaged cells were either directly expanded to larger formats for eventual cloning if they were judged to have appropriate editing or frozen by overlaying and gently mixing equivalent volume 20% mTeSR media with provided 5X supplement (Catalog: 05850, STEMCELL Technologies, Vancouver, BC, Canada) with added 5 mL (1% v/v) P/S (Catalog: 15140-122, Thermo Fisher Scientific, Waltham, MA, USA), 60% Knock-out Serum (Catalog: 10828-028, Thermo Fisher Scientific, Waltham, MA, USA) and 20% DMSO (Catalog: D2650, SigmaAldrich, Inc., St. Louis, MO, USA) at equivalent volume, then transferring into Cryo.S-vials (Catalog: 976561, Greiner Bio-One, Monroe, NC, USA).

### Flow cytometry methods

Cell populations were analyzed for fluorescent protein expression using a CytoFLEX S flow cytometer (Beckman Coulter, San Jose, CA, USA). Single cell suspensions were isolated from each experimental condition 3-5 days after transfection, depending on cell confluency. Cells were Accutase-treated to generate single cell suspensions as previously described, then pelleted in a 96-well U-bottom plate and resuspended in 100-150 μL PBS +1% FBS with DAPI (2 μg/mL). Cells were gated to exclude doublets by FSC-A vs FSC-H followed by the removal of DAPI-positive dead cells. To set positive cell gating, a negative non-targeting AAVS1 transfected, and AAV transduced control cell population was used to set the gate quadrants. For single color experiments, PC5 Area, an unrelated fluorescent parameter, was used to help visualize two-dimensional dot plots and define positive populations. For two-color experiments, mEGFP area and mTagRFP area dot plots were utilized, and quadrant gating was set using the negative control cells as described for single color experiments.

### Droplet digital PCR (ddPCR)

Primers and probes (IDT, Coralville, IA, USA) specific to each knock-in locus were designed with one primer external to the homology arm, the other within the tag sequence, and the probe also within the tag sequence. Primers were prepared for each knock-in on either the 5’ or 3’ side of the predicted knock-in sequence based on our ability to synthesize an accompanying DNA fragment (IDT, Coralville, IA, USA or Twist Biosciences, South San Francisco, CA) for use in positive control assays for ddPCR. Probes were modified with FAM dye and Black Six quencher molecules. Primers were chosen on the side of the donor construct with the least sequence complexity. Amplicon regions were additionally synthesized (Azenta Life Sciences, Seattle, WA USA or Twist Biosciences, South San Francisco, CA) to serve as positive controls. PCR primers were designed to amplify either the 5’ junction (spanning the 5’ homology arm between the distal 5’ genome and the middle of the FP insertion) or the 3’ junction (spanning the 3’ homology arm from the middle of the FP insertion to the 3’ distal genome). Probes were then designed to target the 5’ or 3’ end of the FP. For each ddPCR mix, primers and corresponding junctional probe were mixed with 2X Supermix for probes, no dUTP, HINDIII ((20,000 U/mL) (Catalog: R3104S, New England Biolabs, Ipswich, MA, USA) restriction enzyme, and water for a 1X mixture used to generate droplets (BioRad, Hercules, CA, USA). Appropriate positive (synthetic PCR templates or previously edited clones with known correct junctional sizes) and negative controls (unedited or non-targeting gRNA controls) were used to benchmark assay efficacy. Assays were cycled under the following conditions: 94 °C for 10:00 (min:sec); 94 °C for 0:30; 60 °C for 1:00; 72 °C for 10:00 (repeat steps 2-5 39x); 72 °C for 10:00; 12 °C hold. The ddPCR protocol was modified from previously described (Roberts et al., 2017) to include a 10 mine extension time to reflect the larger target amplicon. Primer, probe and positive control sequences for each target can be found in Table 1. ddPCR droplets were prepared as previously described (Roberts et al., 2017). RPP30 control locus ddPCR reagents were from BioRad (Catalog: 1863024, BioRad, Hercules, CA, USA). Primers and other probes were from IDT and listed in Figure 7.

**Figure 7.**
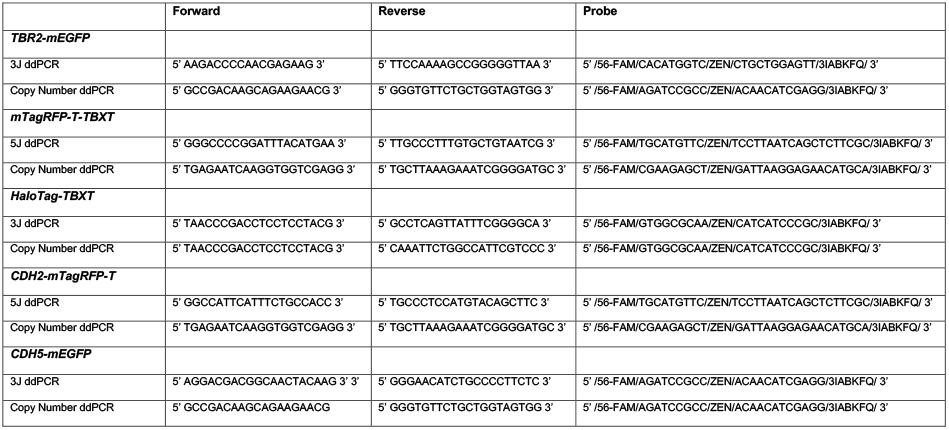
ddPCR primers and probes used in this study.

### Quantifying INDELs via Inference of CRISPR Edits (ICE)

PCR primers (see below) were specifically designed to amplify genomic DNA segments ranging from 500 to 800 base pairs, encompassing the RNP cleavage site. Genomic DNA was extracted from lysed samples using the Quick Extract DNA Extraction Solution (Catalog: QE09050, Bio-Search Technologies, VWR, Radnor, PA, USA), following the protocol provided by the manufacturer. Amplified PCR products were subsequently sequenced by Sanger sequencing, outsourced to Azenta Life Sciences (Seattle, WA, USA). For sequence analysis, the resulting .ab1 files were uploaded to the ICE analysis tool available online at https://ice.synthego.com. The platform required designation of control versus experimental files. In the case of knock-in experiments, only the guide RNA sequence was entered. Additionally, the sequence of the donor DNA, inclusive of both homology arms, was provided. It is imperative that each homology arm aligns with at least 15 base pairs of the target genomic sequence, with an optimal alignment exceeding 50 base pairs. The ICE analysis tool then generated a report for each sample, including:

1. Sample: A unique identifier for each sample.
2. Guide Target: The sequence (17-23 base pairs) to which the gRNA binds within the genomic DNA, not including the Protospacer Adjacent Motif (PAM).
3. PAM Sequence: The specific PAM sequence recognized by the Cas9 nuclease.
4. INDEL Percentage: The fraction of sequences exhibiting INDELs as a proxy for CRISPR/Cas9 editing efficiency. This metric encompasses all unpredicted sequences, indicating knockout events.
5. Model Fit (R^2^): The Pearson correlation coefficient, reflecting the accuracy of the INDEL distribution model in describing the Sanger sequencing data from edited samples.

### PCR validation of isolated clones

In the initial phase of clone screening, a technique called “full allele PCR” was utilized, involving a single PCR reaction with external primers flanking the gene-specific junctions to amplify both modified and unaltered alleles simultaneously. Subsequently, a second round of screening employed PCR to amplify areas encompassing the inserted tag, including the left and right homology arms, the fluorescent protein, linker sequence (if applicable), and adjacent genomic regions, using two sets of overlapping primers specific to the target gene. The final stage of screening involved PCR amplification of both modified and unmodified alleles with primers surrounding the RNP cut site, with sequencing of the PCR products from clones confirmed to possess wild-type untagged alleles to assess for NHEJ.

The PCR mixtures for the reactions described here were composed using PrimeStar (Takara Bio, Shiga, Japan) with a 2X GC buffer, DNTPs at a concentration of 200 μM, 1 unit of PrimeStar HS polymerase, primers at 800 nM, and 10 ng of genomic DNA, all within a total volume of 25 μL. The cycling parameters for the PCR were set as follows: an initial 6 cycles of denaturation at 98 °C for 10 sec-, annealing starting at 70 °C for 5 sec-(decreasing by 2 °C per cycle), and extension at 72 °C for 60 sec-followed by 32 cycles of denaturation at 98 °C for 10 sec-, annealing at 54 °C for 5 sec-, and extension at 72 °C for 60 sec-, with a final hold at 12 °C. Primers listed for all PCRs are listed in Figure 8.

**Figure 8.**
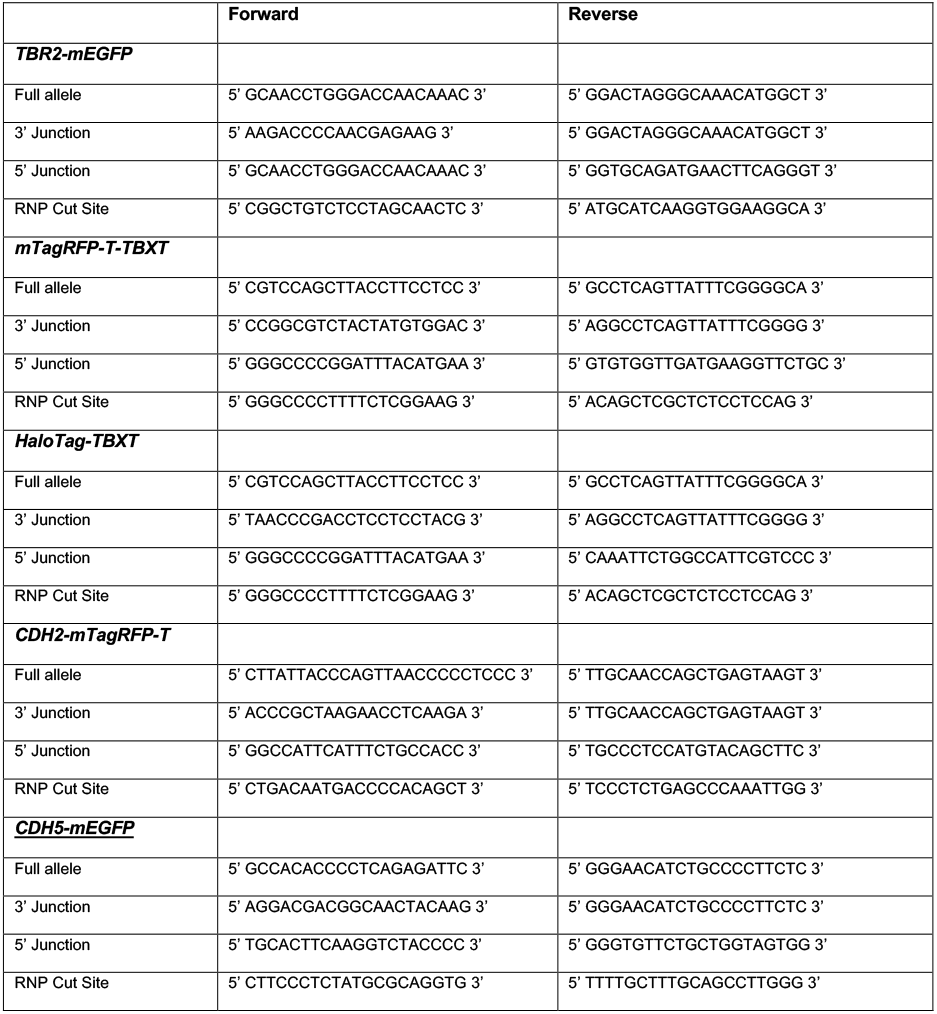
Primers used for clone screening in this study.

## Acknowledgements

We thank Danuta Sastre, Stephanie Q. Dinh, Helen Anderson, Angelique Nelson, Joy Arakaki and Youngmee Sul for assistance with experiments and Renata M. Martin for helpful experimental advice. The WTC-11 hiPSC line used to create our gene-edited cell lines was provided by the Bruce R. Conklin Laboratory at the Gladstone Institute and UCSF. We acknowledge the Allen Institute for Cell Science founder, Paul G. Allen, for his vision, encouragement, and support.

## Supplementary Information

**Figure 1.**
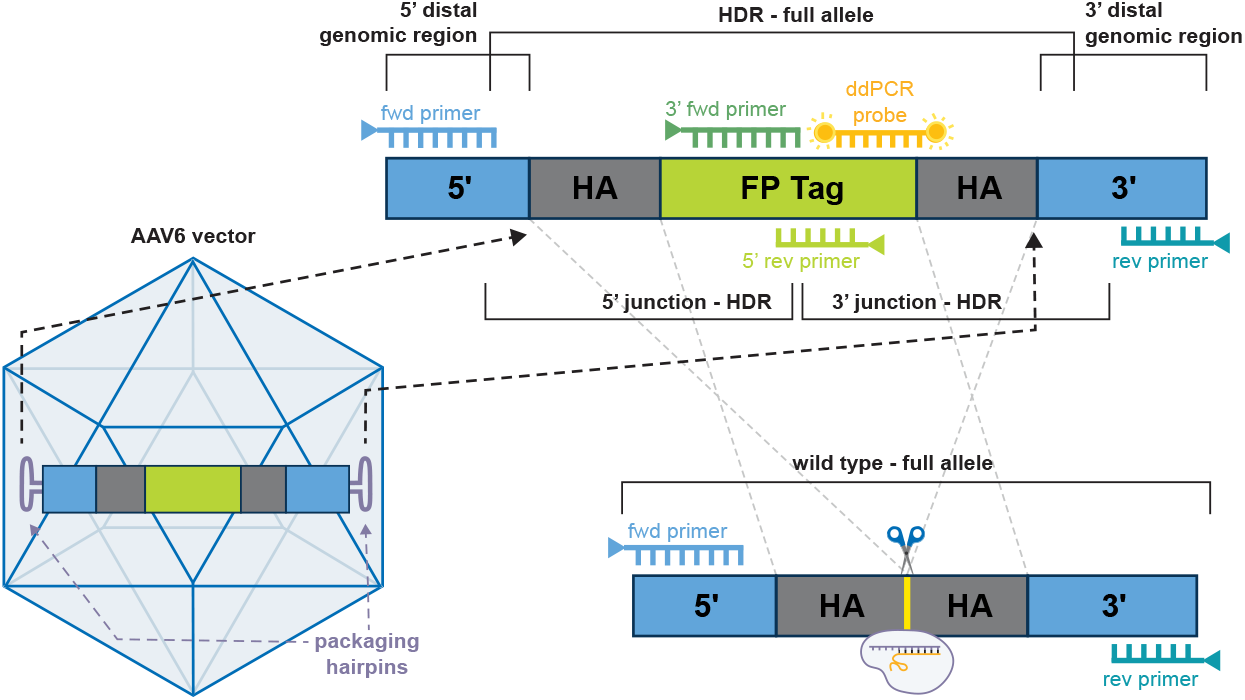
Overview of template and primer design. **(A)** An overview of donor template, primer, and probe design for large knock-ins using the AAV6 vector. Primers are designed outside of the homology arms (HA) (blue) to amplify region around the insertion. Additional primers are made to the fluorescent protein tag (green) to be used in combination with the primers (blue) to amplify 5’ junctions and 3’ junctions. Additionally, ddPCR probes (yellow) are designed internal to the FP insertion. These primers are illustrated showing the ideal HDR-edited full allele on the top and the unedited allele on the bottom. The RNP cut site in the unedited allele is shown with scissors and a yellow line in the bottom graphic.

**Figure 2.**
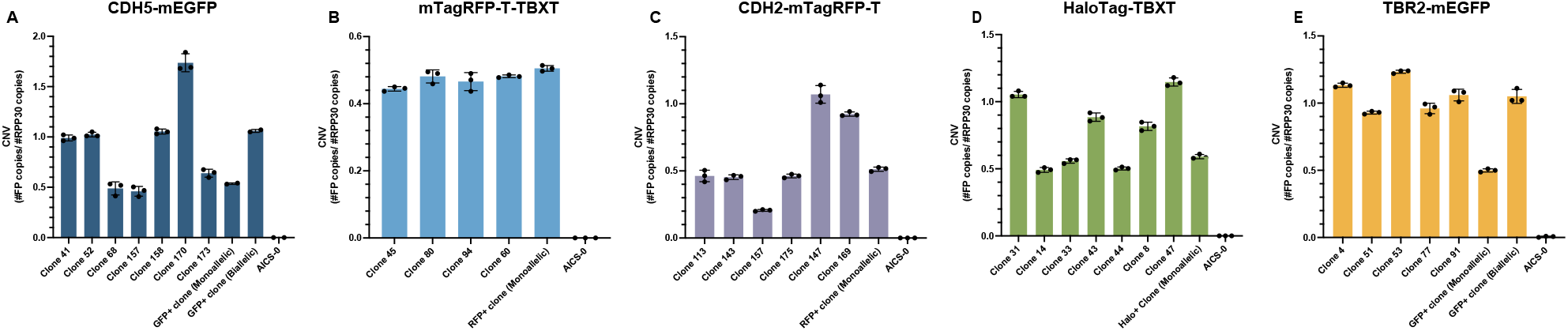
Copy Number Variance (CNV) identification of edited alleles. **(A-E)** CNV shown as the number of FP positive copies over the number of RPP30 copies (reference gene). CNV values around 0.5 are monoallelic and values around 1 are biallelic. Clones surveilled along with positive control (FP+ clone) and negative control (AICS-0/unedited cells) are written on the x-axis.

**Figure 3.**
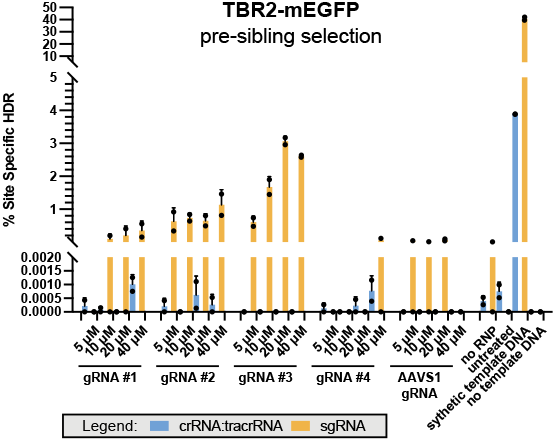
Percent site specific HDR comparing the usage of crRNA:tracrRNA vs. sgRNA. **(A)** *TBR2-mEGFP* editing is compared using two different gRNA methods, crRNA:tracrRNA (blue bars) vs. sgRNA (yellow bars). On the x-axis are gRNA sequences #1-4 with a range of RNP concentrations. On the y-axis is the percentage of site specific HDR. Note the first break in the y-axis is at 0.002%. Generally, sgRNA had higher percentages of HDR compared to crRNA complexes; in addition, HDR percentage increased with RNP concentration.

## References

Artegiani, B., Hendriks, D., Beumer, J., Kok, R., Zheng, X., Joore, I., Chuva De Sousa Lopes, S., Van Zon, J., Tans, S., and Clevers, H. Fast and efficient generation of knock-in human organoids using homology-independent CRISPR–Cas9 precision genome editing. Nature Cell Biology, 22(3):321–331, Mar. 2020. doi: 10.1038/s41556-020-0472-5.

Bae, S., Kweon, J., Kim, H. S., and Kim, J.-S. Microhomology-based choice of Cas9 nuclease target sites. Nature Methods, 11(7):705–706, July 2014. doi: 10.1038/nmeth.3015.

Bak, R. O., Dever, D. P., Reinisch, A., Cruz Hernandez, D., Majeti, R., and Porteus, M. H. Multiplexed genetic engineering of human hematopoietic stem and progenitor cells using CRISPR/Cas9 and AAV6. eLife, 6:e27873, Sept. 2017. doi: 10.7554/eLife.27873.

Boyle, E. A., Becker, W. R., Bai, H. B., Chen, J. S., Doudna, J. A., and Greenleaf, W. J. Quantification of Cas9 binding and cleavage across diverse guide sequences maps landscapes of target engagement. Science Advances, 7 (8):eabe5496, Feb. 2021. doi: 10.1126/sciadv.abe5496.

Bukhari, H. and Müller, T. Endogenous Fluorescence Tagging by CRISPR. Trends in Cell Biology, 29(11):912–928, Nov. 2019. doi: 10.1016/j.tcb.2019.08.004.

Charlesworth, C. T., Camarena, J., Cromer, M. K., Vaidyanathan, S., Bak, R. O., Carte, J. M., Potter, J., Dever, D. P., and Porteus, M. H. Priming Human Repopulating Hematopoietic Stem and Progenitor Cells for Cas9/sgRNA Gene Targeting. Molecular Therapy - Nucleic Acids, 12:89–104, Sept. 2018. doi: 10.1016/j.omtn.2018.04.017.

Cho, N. H., Cheveralls, K. C., Brunner, A.-D., Kim, K., Michaelis, A. C., Raghavan, P., Kobayashi, H., Savy, L., Li, J. Y., Canaj, H., Kim, J. Y. S., Stewart, E. M., Gnann, C., McCarthy, F., Cabrera, J. P., Brunetti, R. M., Chhun, B. B., Dingle, G., Hein, M. Y., Huang, B., Mehta, S. B., Weissman, J. S., Gómez-Sjöberg, R., Itzhak, D. N., Royer, L. A., Mann, M., and Leonetti, M. D. OpenCell: Endogenous tagging for the cartography of human cellular organization. Science, 375(6585):eabi6983, Mar. 2022. doi: 10.1126/science.abi6983.

Conant, D., Hsiau, T., Rossi, N., Oki, J., Maures, T., Waite, K., Yang, J., Joshi, S., Kelso, R., Holden, K., Enzmann, B. L., and Stoner, R. Inference of CRISPR Edits from Sanger Trace Data. The CRISPR Journal, 5(1):123–130, Feb. 2022. doi: 10.1089/crispr.2021.0113.

Dambournet, D., Sochacki, K. A., Cheng, A. T., Akamatsu, M., Taraska, J. W., Hockemeyer, D., and Drubin, D. G. Genome-edited human stem cells expressing fluorescently labeled endocytic markers allow quantitative analysis of clathrin-mediated endocytosis during differentiation. Journal of Cell Biology, 217(9):3301–3311, Sept. 2018. doi: 10.1083/jcb.201710084.

Dang, Y., Jia, G., Choi, J., Ma, H., Anaya, E., Ye, C., Shankar, P., and Wu, H. Optimizing sgRNA structure to improve CRISPR-Cas9 knock-out efficiency. Genome Biology, 16(1):280, Dec. 2015. doi: 10.1186/s13059-015-0846-3.

Doench, J. G., Fusi, N., Sullender, M., Hegde, M., Vaimberg, E. W., Donovan, K. F., Smith, I., Tothova, Z., Wilen, C., Orchard, R., Virgin, H. W., Listgarten, J., and Root, D. E. Optimized sgRNA design to maximize activity and minimize off-target effects of CRISPR-Cas9. Nature Biotechnology, 34(2):184–191, Feb. 2016. doi: 10.1038/nbt.3437.

Fehér, A., Schnúr, A., Muenthaisong, S., Bellák, T., Ayaydin, F., Várady, G., Kemter, E., Wolf, E., and Dinnyés, A. Establishment and characterization of a novel human induced pluripotent stem cell line stably expressing the iRFP720 reporter. Scientific Reports, 12(1):9874, June 2022. doi: 10.1038/s41598-022-12956-1.

Fu, Y.-W., Dai, X.-Y., Wang, W.-T., Yang, Z.-X., Zhao, J.-J., Zhang, J.-P., Wen, W., Zhang, F., Oberg, K. C., Zhang, L., Cheng, T., and Zhang, X.-B. Dynamics and competition of CRISPR–Cas9 ribonucleoproteins and AAV donor-mediated NHEJ, MMEJ and HDR editing. Nucleic Acids Research, 49(2): 969–985, Jan. 2021. doi: 10.1093/nar/gkaa1251.

Gao, Y., Hisey, E., Bradshaw, T. W., Erata, E., Brown, W. E., Courtland, J. L., Uezu, A., Xiang, Y., Diao, Y., and Soderling, S. H. Plug-and-Play Protein Modification Using Homology-Independent Universal Genome Engineering. Neuron, 103(4):583–597.e8, Aug. 2019. doi: 10.1016/j.neuron.2019.05.047.

Gerbin, K. A., Grancharova, T., Donovan-Maiye, R. M., Hendershott, M. C., Anderson, H. G., Brown, J. M., Chen, J., Dinh, S. Q., Gehring, J. L., Johnson, G. R., Lee, H., Nath, A., Nelson, A. M., Sluzewski, M. F., Viana, M. P., Yan, C., Zaunbrecher, R. J., Cordes Metzler, K. R., Gaudreault, N., Knijnenburg, T. A., Rafelski, S. M., Theriot, J. A., and Gunawardane, R. N. Cell states beyond transcriptomics: Integrating structural organization and gene expression in hiPSC-derived cardiomyocytes. Cell Systems, 12(6):670–687.e10, June 2021. doi: 10.1016/j.cels.2021.05.001.

Kreitzer, F. R., Salomonis, N., Sheehan, A., Huang, M., Park, J. S., Spindler, M. J., Lizarraga, P., Weiss, W. A., So, P.-L., and Conklin, B. R. A robust method to derive functional neural crest cells from human pluripotent stem cells. American Journal of Stem Cells, 2(2):119–131, 2013.

Kurokawa, Y. K., Yin, R. T., Shang, M. R., Shirure, V. S., Moya, M. L., and George, S. C. Human Induced Pluripotent Stem Cell-Derived Endothelial Cells for Three-Dimensional Microphysiological Systems. Tissue Engineering Part C: Methods, 23(8):474–484, Aug. 2017. doi: 10.1089/ten.tec.2017.0133.

Lasham, A., Tsai, P., Fitzgerald, S. J., Mehta, S. Y., Knowlton, N. S., Braithwaite, A. W., and Print, C. G. Accessing a New Dimension in TP53 Biology: Multiplex Long Amplicon Digital PCR to Specifically Detect and Quantitate Individual TP53 Transcripts. Cancers, 12(3):769, Mar. 2020. doi: 10.3390/cancers12030769.

Lau, C.-H., Tin, C., and Suh, Y. CRISPR-based strategies for targeted transgene knock-in and gene correction. Faculty Reviews, 9, Dec. 2020. doi: 10.12703/r/9-20.

Martin, R. M., Ikeda, K., Cromer, M. K., Uchida, N., Nishimura, T., Romano, R., Tong, A. J., Lemgart, V. T., Camarena, J., Pavel-Dinu, M., Sindhu, C., Wiebking, V., Vaidyanathan, S., Dever, D. P., Bak, R. O., Laustsen, A., Lesch, B. J., Jakobsen, M. R., Sebastiano, V., Nakauchi, H., and Porteus, M. H. Highly Efficient and Marker-free Genome Editing of Human Pluripotent Stem Cells by CRISPR-Cas9 RNP and AAV6 Donor-Mediated Homologous Recombination. Cell Stem Cell, 24(5):821–828.e5, May 2019. doi: 10.1016/j.stem.2019.04.001.

Mikkelsen, N. S. and Bak, R. O. Enrichment strategies to enhance genome editing. Journal of Biomedical Science, 30(1):51, July 2023. doi: 10.1186/s12929-023-00943-1.

Miyaoka, Y., Chan, A. H., Judge, L. M., Yoo, J., Huang, M., Nguyen, T. D., Lizarraga, P. P., So, P.-L., and Conklin, B. R. Isolation of single-base genome-edited human iPS cells without antibiotic selection. Nature Methods, 11(3): 291–293, Mar. 2014. doi: 10.1038/nmeth.2840.

Miyaoka, Y., Chan, A. H., and Conklin, B. R. Using Digital Polymerase Chain Reaction to Detect Single-Nucleotide Substitutions Induced by Genome Editing. Cold Spring Harbor Protocols, 2016(8):pdb.prot086801, Aug. 2016. doi: 10.1101/pdb.prot086801.

Rafelski, S. M. and Theriot, J. A. Establishing a conceptual framework for holistic cell states and state transitions. Cell, 187(11):2633–2651, May 2024. doi: 10.1016/j.cell.2024.04.035.

Riesenberg, S., Kanis, P., Macak, D., Wollny, D., Düsterhöft, D., Kowalewski, J., Helmbrecht, N., Maricic, T., and Pääbo, S. Efficient high-precision homology-directed repair-dependent genome editing by HDRobust. Nature Methods, 20(9):1388–1399, Sept. 2023. doi: 10.1038/s41592-023-01949-1.

Roberts, B., Haupt, A., Tucker, A., Grancharova, T., Arakaki, J., Fuqua, M. A., Nelson, A., Hookway, C., Ludmann, S. A., Mueller, I. A., Yang, R., Horwitz, R., Rafelski, S. M., and Gunawardane, R. N. Systematic gene tagging using CRISPR/Cas9 in human stem cells to illuminate cell organization. Molecular Biology of the Cell, 28(21):2854–2874, Oct. 2017. doi: 10.1091/mbc.e17-03-0209.

Roberts, B., Hendershott, M. C., Arakaki, J., Gerbin, K. A., Malik, H., Nelson, A., Gehring, J., Hookway, C., Ludmann, S. A., Yang, R., Haupt, A., Grancharova, T., Valencia, V., Fuqua, M. A., Tucker, A., Rafelski, S. M., and Gunawardane, R. N. Fluorescent Gene Tagging of Transcriptionally Silent Genes in hiPSCs. Stem Cell Reports, 12(5):1145–1158, May 2019. doi: 10.1016/j.stemcr.2019.03.001.

Schubert, M. S., Thommandru, B., Woodley, J., Turk, R., Yan, S., Kurgan, G., McNeill, M. S., and Rettig, G. R. Optimized design parameters for CRISPR Cas9 and Cas12a homology-directed repair. Scientific Reports, 11(1):19482, Sept. 2021. doi: 10.1038/s41598-021-98965-y.

Sheppard, H. E., Dall’Agnese, A., Park, W. D., Shamim, M. H., Dubrulle, J., Johnson, H. L., Stossi, F., Cogswell, P., Sommer, J., Levy, J., Sharifnia, T., Wawer, M. J., Nabet, B., Gray, N. S., Clemons, P. A., Schreiber, S. L., Workman, P., Young, R. A., and Lin, C. Y. Targeted brachyury degradation disrupts a highly specific autoregulatory program controlling chordoma cell identity. Cell Reports Medicine, 2(1):100188, Jan. 2021. doi: 10.1016/j.xcrm.2020.100188.

Simkin, D., Papakis, V., Bustos, B. I., Ambrosi, C. M., Ryan, S. J., Baru, V., Williams, L. A., Dempsey, G. T., McManus, O. B., Landers, J. E., Lubbe, S. J., George, A. L., and Kiskinis, E. Homozygous might be hemizygous: CRISPR/Cas9 editing in iPSCs results in detrimental on-target defects that escape standard quality controls. Stem Cell Reports, 17(4):993–1008, Apr. 2022. doi: 10.1016/j.stemcr.2022.02.008.

Skarnes, W. C., Pellegrino, E., and McDonough, J. A. Improving homology-directed repair efficiency in human stem cells. Methods, 164-165:18–28, July 2019. doi: 10.1016/j.ymeth.2019.06.016.

Soldner, F., Laganière, J., Cheng, A. W., Hockemeyer, D., Gao, Q., Alagappan, R., Khurana, V., Golbe, L. I., Myers, R. H., Lindquist, S., Zhang, L., Guschin, D., Fong, L. K., Vu, B. J., Meng, X., Urnov, F. D., Rebar, E. J., Gregory, P. D., Zhang, H. S., and Jaenisch, R. Generation of Isogenic Pluripotent Stem Cells Differing Exclusively at Two Early Onset Parkinson Point Mutations. Cell, 146(2):318–331, July 2011. doi: 10.1016/j.cell.2011.06.019.

Suchy, F. P., Karigane, D., Nakauchi, Y., Higuchi, M., Zhang, J., Pekrun, K., Hsu, I., Fan, A. C., Nishimura, T., Charlesworth, C. T., Bhadury, J., Nishimura, T., Wilkinson, A. C., Kay, M. A., Majeti, R., and Nakauchi, H. Genome engineering with Cas9 and AAV repair templates generates frequent concatemeric insertions of viral vectors. Nature Biotechnology, Apr. 2024. doi: 10.1038/s41587-024-02171-w.

Tosic, J., Kim, G.-J., Pavlovic, M., Schröder, C. M., Mersiowsky, S.-L., Barg, M., Hofherr, A., Probst, S., Köttgen, M., Hein, L., and Arnold, S. J. Eomes and Brachyury control pluripotency exit and germ-layer segregation by changing the chromatin state. Nature Cell Biology, 21(12):1518–1531, Dec. 2019. doi: 10.1038/s41556-019-0423-1.

Viana, M. P., Chen, J., Knijnenburg, T. A., Vasan, R., Yan, C., Arakaki, J. E., Bailey, M., Berry, B., Borensztejn, A., Brown, E. M., Carlson, S., Cass, J. A., Chaudhuri, B., Cordes Metzler, K. R., Coston, M. E., Crabtree, Z. J., Davidson, S., DeLizo, C. M., Dhaka, S., Dinh, S. Q., Do, T. P., Domingus, J., Donovan-Maiye, R. M., Ferrante, A. J., Foster, T. J., Frick, C. L., Fujioka, G., Fuqua, M. A., Gehring, J. L., Gerbin, K. A., Grancharova, T., Gregor, B. W., Harrylock, L. J., Haupt, A., Hendershott, M. C., Hookway, C., Horwitz, A. R., Hughes, H. C., Isaac, E. J., Johnson, G. R., Kim, B., Leonard, A. N., Leung, W. W., Lucas, J. J., Ludmann, S. A., Lyons, B. M., Malik, H., McGregor, R., Medrash, G. E., Meharry, S. L., Mitcham, K., Mueller, I. A., Murphy-Stevens, T. L., Nath, A., Nelson, A. M., Oluoch, S. A., Paleologu, L., Popiel, T. A., Riel-Mehan, M. M., Roberts, B., Schaefbauer, L. M., Schwarzl, M., Sherman, J., Slaton, S., Sluzewski, M. F., Smith, J. E., Sul, Y., Swain-Bowden, M. J., Tang, W. J., Thirstrup, D. J., Toloudis, D. M., Tucker, A. P., Valencia, V., Wiegraebe, W., Wijeratna, T., Yang, R., Zaunbrecher, R. J., Labitigan, R. L. D., Sanborn, A. L., Johnson, G. T., Gunawardane, R. N., Gaudreault, N., Theriot, J. A., and Rafelski, S. M. Integrated intracellular organization and its variations in human iPS cells. Nature, 613(7943):345–354, Jan. 2023. doi: 10.1038/s41586-022-05563-7. Number: 7943 Publisher: Nature Publishing Group.

Zhang, J.-P., Li, X.-L., Li, G.-H., Chen, W., Arakaki, C., Botimer, G. D., Baylink, D., Zhang, L., Wen, W., Fu, Y.-W., Xu, J., Chun, N., Yuan, W., Cheng, T., and Zhang, X.-B. Efficient precise knockin with a double cut HDR donor after CRISPR/Cas9-mediated double-stranded DNA cleavage. Genome Biology, 18(1):35, Dec. 2017. doi: 10.1186/s13059-017-1164-8.

